# Experimental evidence of non-classical brain functions

**DOI:** 10.1101/219931

**Authors:** Christian Kerskens, David López Pérez

## Abstract

Recent proposals in quantum gravity have suggested that unknown systems can mediate entanglement between two known quantum systems, if and only if the mediator itself is non-classical. This approach may be applicable to the brain, where speculations about quantum operations in consciousness and cognition have a long history.

Proton spins of bulk water, which most likely interfere with any brain function, can act as the known quantum systems. If an unknown mediator exists, then NMR methods based on multiple quantum coherence (MQC) can act as entanglement witness. However, there are doubts that today’s NMR signals can contain quantum correlations in general, and specifically in the brain environment. Here, we used a witness protocol based on zero quantum coherence (ZQC) whereby we minimised the classical signals to circumvent the NMR detection limits for quantum correlation. For short repetitive periods, we found evoked signals in most parts of the brain, whereby the temporal appearance resembled heartbeat-evoked potentials (HEPs). We found that those signals had no correlates with any classical NMR contrast. Similar to HEPs, the evoked signal depended on conscious awareness. Consciousness-related or electrophysiological signals are unknown in NMR. Remarkably, these signals only appeared if the local properties of the magnetisation were reduced. Our findings suggest that we may have witnessed entanglement mediated by consciousness-related brain functions. Those brain functions must then operate non-classically, which would mean that consciousness is non-classical.

## I. INTRODUCTION

Quantum mechanisms are at work in sensory systems feeding the brain with information [1–3]. Foremost in magneto-reception [4], there is no doubt that only quantum mechanical effects can explain its sensitivity [3]. It has been suggested that entangled radical electron pairs are involved.

Beyond those sensory inputs, more complex brain functionalities depend on the presence of specific nuclear spins. For example, Lithium-6 isotopes with nuclear spin 1 increase activity of complex behaviour in contrast to Lithium-7 isotopes with 3/2 spin where it decreases [5]. Similar, Xenon isotopes with 1/2 spin are effective anaesthetizers in contrast to Xenon isotopes with spin 0 which have only little effects [6].

However, nuclear spins can, like electron spins, influence chemical reactions [7], which then lead to macroscopic results as commonly observed in physiology. Whether those or other macroscopic systems in the brain can be non-classical, is still unknown. Experimental methods, which could distinguish classical from quantum correlations in the living brain, haven’t yet been established. In this respect, recent proposals in quantum gravity [8, 9] may help to overcome experimental restrictions in living systems. Those proposals use auxiliary quantum systems for which they showed that if a system can mediate entanglement between auxiliary quantum systems then the mediator itself is non-classical.

If a cerebral mediator of this kind exist, then it is likely that the entanglement plays an important role in the brain. Although, quantum computing can be achieved without entanglement [10], it is commonly believed that entanglement is essential to play out its full advantages [10]. Therefore, it is likely that entanglement, if medi-ated by any brain function at all, may only occur during brain activity.

Hence, the experimental demands on an auxiliary quantum system are that they can be measured non-invasively in the conscious-aware brain, and further that entanglement can be witnessed.

A non-invasive approach offers NMR. The nuclear spins are quantum systems which could, in theory, be entangled by a cerebral mediator. NMR sequences based on multiple quantum coherence (MQC) are also able to witness entanglement [11].

The MQC entanglement witness relies on bounds which, for applications in biology, may be based on the maximal classical signal achievable. The maximal classical MQC signal in fluids have been estimated on the basis of the intermolecular MQC (iMQC) approach [12]. The iMQC signal, despite the naming, is an entirely classical signal because it can also be the classically derived [13] which is known as multiple spin echo (MSE)[14, 15]. Therefore, it can be used as the classical bound.

Further, an exclusion of classicality can also be argued on the following basis. A single quantum coherence (SQC) which is weighted by susceptibility or diffusion contrast may respond similar to physiological changes as the iMQC contrast which is caused by long-range dependency or rotational symmetry breaking [16–19]. Hence, a signal change in a MQC sequence with no corresponding diffusion or 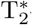-weighted SQC signal is most likely non-classically. With this knowledge at hand, we can now search for situations in which witnessing entanglement may be possible. As mentioned before brain activity, or more concretely brain computation, may play a crucial role in the creation of cerebral entanglement. Hence, we can make additional observations specific to the brain. We propose the following conditions:

1. Sufficient condition for witness – Non-invasive direct detection of brain computation is possible using electrophysiological measurements. With today’s technology, corresponding MRI signals are undetectable. Therefore we conclude that a detection of an electrophysiological event with a conventional MRI system, which classically is not possible, would be sufficient to witness entanglement.
2. Necessary condition for witness – The brain is able to operate without any external magnetic fields which means that, without a brain function at work, all states are initially mixed. Hence, the assumed brain function producing entanglement, must use a kind of quantum distillation process [20] on mixed states [21]. Therefore, we conclude that the NMR signal must initially be saturated.

The following two arguments underpin the importance of saturation for the detection further. The unusual dispense of (pseudo-)pure states on which the MR signal is normally constituted, circumnavigates the major problem that entanglement of pure spins, which are in close proximity, is highly unlikely [22]. Further, the saturation of pure local states may serve the existence of non-localities because local and non-local properties can be complimentary [23, 24].

Now, we are in the position to address the question whether the brain can mediate entanglement, experimentally. Based on the above considerations, we explored if the conscious-aware brain may use entanglement during computing. As indicators of brain computation, we focussed on electrophysiological brain waves, which can be observed in the conscious-aware brain at rest.

We acquired MRI time series which were highly saturated and which were able to detect ZQC. Based on the maximal temporal resolution of our method (*<* 5Hz), we focussed on Heartbeat Evoked Potentials (HEPs) [25] which like other electrophysiological signals are far below the detection threshold of conventional MRI sequences.

## II. RESULTS

We used the EPI time series (as described in section III) in human volunteers at rest. The beginning of the sequential RF-pulses train of the EPI time series were used to saturate the magnetisation of the imaging slice. The desired reductions of the local NMR component were normally reached shortly before the equilibrium magnetisation.

Then, we found regular, repeating signal bursts of predominant signal alternations in single volumes of the brain slices as shown in Fig. 1, where the signal peaks of the bursts increased by up to 15 %. In most cases, the alteration was sequential from one image acquisition to the next.

**Figure 1.**
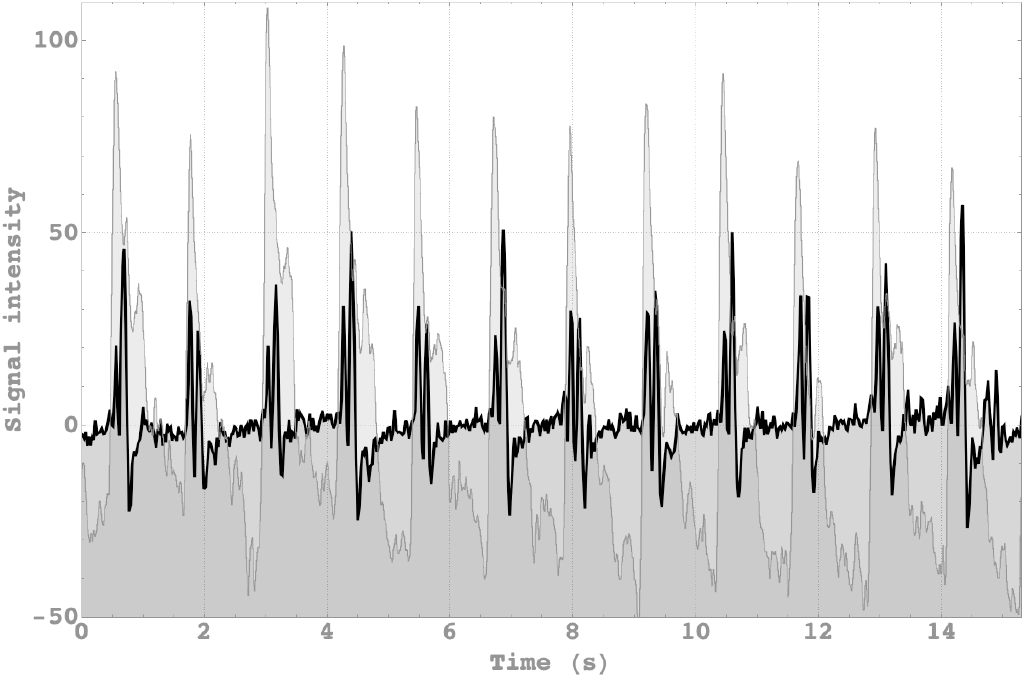
MRI signal time course (Black) during 12 heart cycles compared with simultaneous oximeter reading of a finger (Grey).

In the following, we will focus on the NMR contrast mechanism of the signal first, and then how it related to physiology and mind.

### A. NMR contrast

The burst signal had the following properties;

The burst signal alternated during burst which confirmed that at least two RF pulses were necessary to generate the signal. The two RF-pulses always enwrap an asymmetrical gradient interval *G*_*a*_*T*_*a*_ (Fig. 5), which is the basic pulse design to measure ZQC. The consequential long-range ZQC contrast was verified further by altering sequence parameters.

For rotating the asymmetric gradients *G*_*a*_, we found the characteristic angulation dependency of the dipole-dipole interaction as shown is (Fig. 2A). The plot represents the fitted function | (3 *· cos*^2^[*ϕ*] *−* 1) | (adjusted R^2^ test of goodness-of-fit resulted in R^2^=0.9958) where *ϕ* takes the additional gradients in read and phase direction into account. At the magic angle, the burst signals disappeared. For the flip angle variation, we found the predicted signal course for the ZQC flip angle dependency [26] which was fitted to the data (R^2^=0.9964). Predicted maximum at 45° could be confirmed (Fig 2B). In contrast, the Ernst-angle [27] which is a good indication for the optimum angle for SQC is around 13° (for T_1_ = 1.5s).

**Figure 2.**
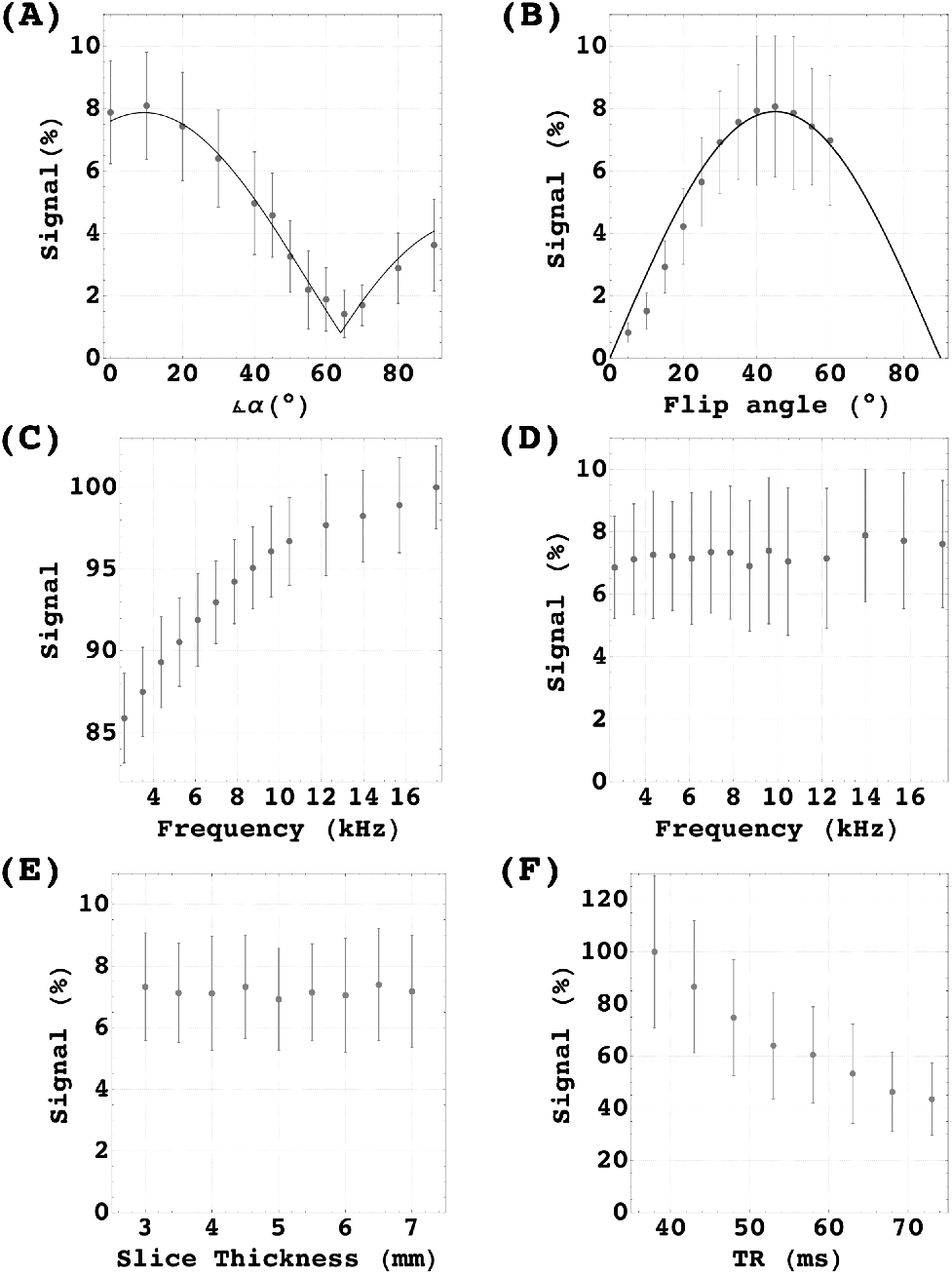
Variation of sequence parameters. Data shows signal averaged over 5 subjects. Error bars represent the standard deviation from the mean. **(A)** Signal intensity plotted against the slice gradient angulation *α* in respect to the magnetic main field. **(B)** Signal plotted against flip angle variation. ZQC prediction plotted in Black. **(C)** Signal intensity plotted against the frequency offset of the saturation slices of the BS and **(D)** averaged signal of the AMP. **(E)** Relative signal change plotted against slice thickness. **(F)** Signal plotted against repetition time.

For the alteration of the off-resonance frequency, we found a typical magnetisation transfer contrast (MTC) change for the baseline which depended on the off-resonance frequency (Fig. 2C). In contrast, the signal intensity showed the remarkable immunity to MTC as expected for ZQC [28] with no significant changes in the same frequency range (Fig. 2D).

The effects of the competing effects, the build up of the ZQC on the one hand and de-phasing over time on the other hand, were studied varying the TR. We found that from 38 ms onwards the signals showed no growth of ZQC. The free induction dominated.

Finally, we varied the slice thickness to study Time-of-flight effects. We found no significant influence on the relative signal.

### B. Physiology and Mind

The periods of signal bursts repeated with the same rate as the heart-beat. We used three temporal reference systems; (a) a finger pulse oximetry, (b) an electrocardiogram (ECG), and (c) the time-of-flight signal of a voxel placed in the superior sagittal sinus. The signal bursts appeared with the pulse from the finger pulse oximetry (Fig. 1). In relation to the ECG, we found using the Cross-Recurrence Quantification Analysis that the maximum burst signal was delayed by 0.3s on average. With the start of the venous outflow, the bursts always ended as shown in Fig. 4 and Fig. 8.

**Figure 3.**
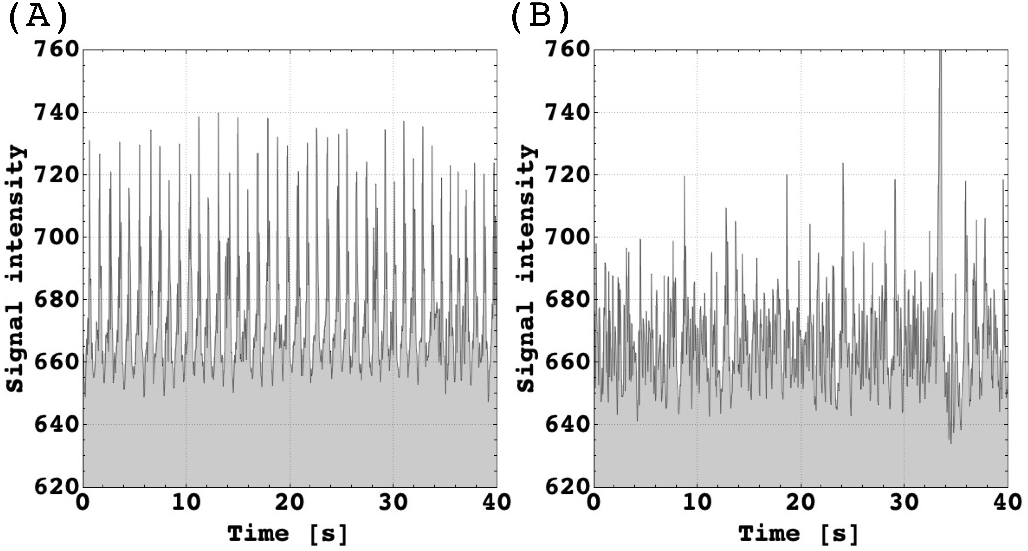
Pattern observed in participant who had reported falling asleep. **(A)** Wake period. **(B)** Asleep, ZQC burst signals declined coincident with an increase of the S/N level. At 34 s, the peak resulted from short head movement

**Figure 4.**
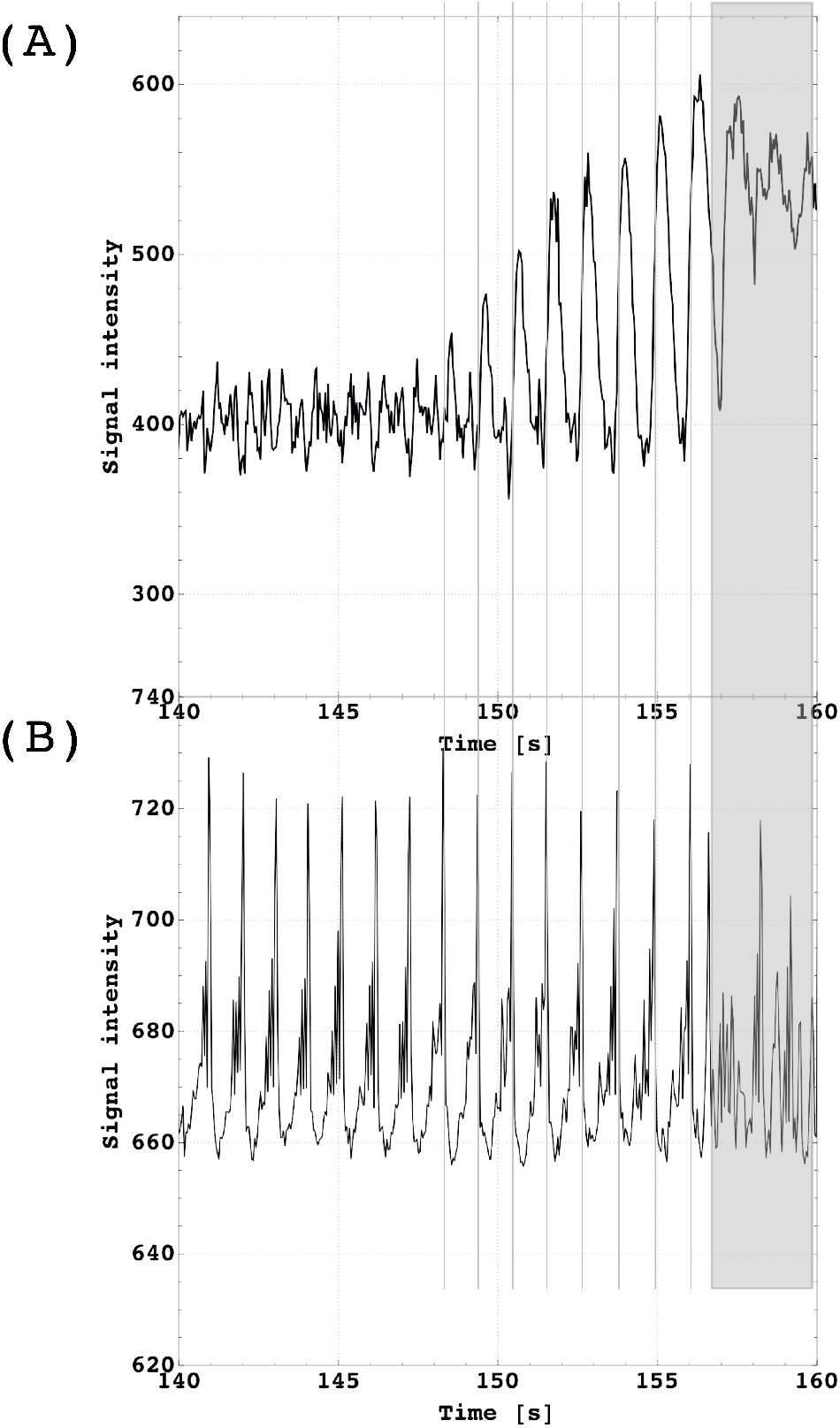
**(A)** Signal time course of an imaging voxel located next to the superior sagittal sinus demonstrates the blood flow increase in response to the CO_2_ challenge (breath-holding). In contrast to the vein signal, the corresponding ZQC signals **(B)** showed no response to CO_2_ activity. Breath-holding started at 140s. Volunteers were instructed to reduce any movement as long as possible (here until at 157s). From 157s, the signal breakdown was subject to movement.

Regarding the duration of the bursts under normal conditions, we mostly observed two sequential peaks which equaled 4 TRs adding up to a time period of 180ms. We also saw longer periods building up to 10 TRs (see Fig. 4B) extending the period to 450ms.

We located the bursts in brain tissue of all slices except around the periventricular area (probably due to movement induced by ventricular pulsation in those regions [29]) as illustrated in Fig. 7.

The global aspect conformed with another interesting feature; the signal could be restored while being averaged over the entire tissue component of the imaging slice (Fig. 1 and Fig. 4B, single voxel time course are shown in Fig. 6).

We also found that the signal did not respond to the CO_2_ challenge (Fig. 4B) in contrast to the SQC signal from the voxel including the superior sagittal sinus (Fig. 4A) which indicated the blood flow response.

During our studies, we also realised that the signal depended on awareness and awakening. In seven participants from whom two had reported to have fallen asleep, we found that the signal pattern declined as shown in Fig. 3. For the final data acquisition, all participants had been asked to stay awake during the imaging protocol. At this point, we no longer detected a sleep pattern. In a case study, we observed the pattern change over a period of 20 minutes which showed a gradual transition from awake to asleep as shown in the appendix at Fig. 9. We used Recurrence Quantification Analysis and Multi-fractal Detrended Fluctuation Analysis to illustrate the difference between wakefulness and the slow decline during the falling asleep period. The analysis shows that periodicity, large and small fluctuations, repeating patterns and their predictability, and the stability of the system were changing over the observation period (Fig. 10).

## III. METHODS

We studied 40 subjects (between 18 and 46 years old) using a 3 Tesla whole-body MRI scanner (Philips, The Netherlands) which was operated with a 32-channel array receiver coil.

Imaging protocols were approved by Trinity College Dublin School of Medicine Research Ethics Committee. All participants of final data acquisition were asked to stay awake and stay still during the imaging protocol, or to report any failure to do so.

Fast gradient-echo EPI (GE-EPI) time series were carried out which had been optimised over a wide range of participants. The finalised parameters were as follows: FA = 45°, TR = 45 ms, TE = 5, voxel size = 3.5 × 3.5 × 3.5 mm, matrix size = 64x64, SENSE factor = 3, bandwidth readout direction = 2148 Hz, saturation pulse thickness/distance = 5/20mm.

Two saturation pulses placed parallel to the imaging slice (Fig. 5) were added which allowed us to vary long-range correlation of the ZSE and MTC. Saturation gradients had a time integral (length x strength) of GT_*s*_ = 5.1 ms x 6.25 mT/m, the crusher gradients in read and slice direction of GT_*c*_ = 1.3 ms x 25 mT/m, the slice rephase gradient of GT_*r*_ = 0.65 ms x 25 mT/m, and the slice termination gradient of GT_*t*_ = 0.65 ms x 15 mT/m. Gradients timing and arrangements are shown in Fig. 5. Gradients relevant for ZSE are shown in the asymmetry field and are marked with indices t, c, r, and s for identification. We rotated the asymmetric gradients in respect to the magnet field starting from coronal 0° to axial 90°in twelve steps; slice angulation *α* related to the angulation from the spin-spin interaction as *φ* = *α−* tan^−1^ ([GT_*c*_*−* GT_*r*_]*/*[2*·* GT_*s*_ + GT_*c*_ + GT_*t*_]) = *α−* 9.6°. Further, we varied the correlation distance via altering the amplitude and the duration of the saturation gradients.

**Figure 5.**
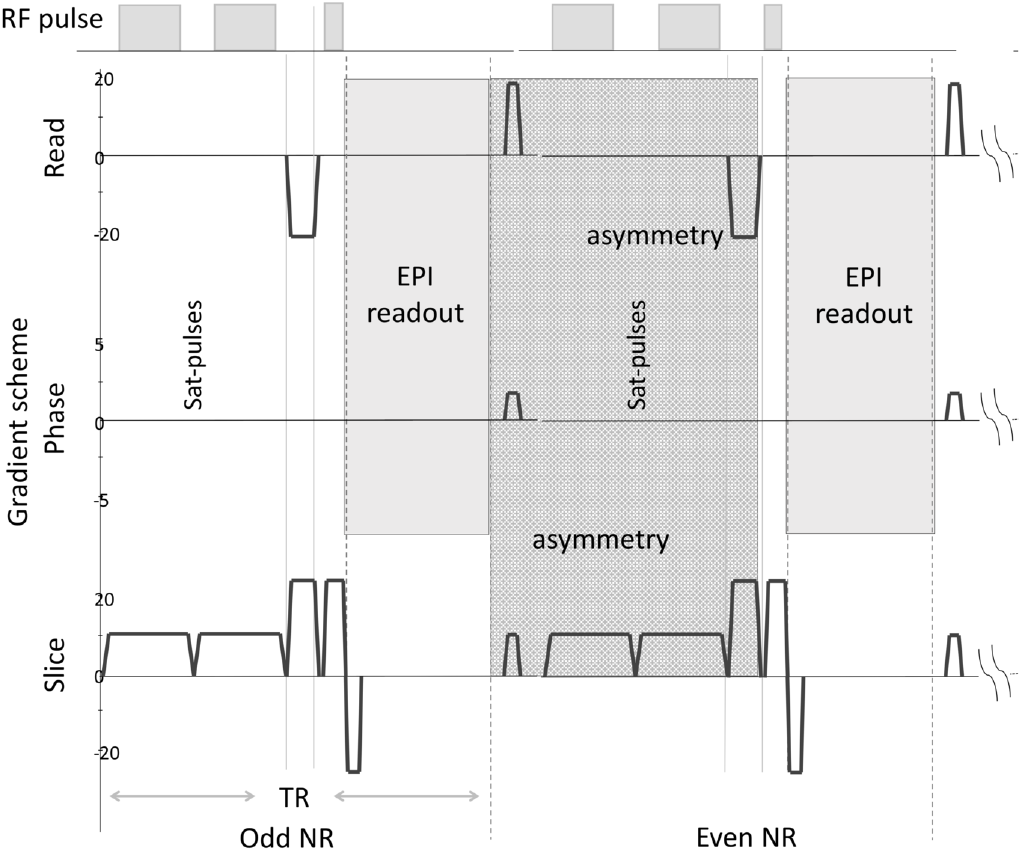
Radio frequency (RF) and Gradient scheme of two consecutive EPI acquisition. The “asymmetry” field includes all asymmetric gradients involved in the ZSE.

We also altered the following sequence parameters in pseudo-randomised orders:

a. variation of the flip angle from 5° to 60° in steps of 5°(60° was the power limit by the specific absorption rate (SAR)).
b. the off-resonance frequency was varied as [2.62, 3.49, 4.36, 5.23, 6.11, 6.98, 7.84, 8.73, 9.60, 10.47, 12.22, 13.96, 15.71, 17.45] kHz.
c. slice thickness from 3 mm to 7 mm in steps of 0.5 mm.
d. repetition time (TR) varied from 38 ms to 73 ms in steps of 5 ms.

Further, we explored the signal distribution over the entire brain. 9 slices (in 5 volunteers) were acquired at different positions, with slices from bottom to the top covering all anatomical regions.

In a breath-holding challenge, four participants were asked to stop breathing for 20 s without taking a deep breath. Body movements were reduced through multiple cushions immobilizing the head.

For the time reference analysis, we used Cross-Recurrence Quantification Analysis [30] to calculate the delay between the R-wave in electrocardiogram (ECG) and the MRI signal. For the calculation, we used the CRP Toolbox [31, 32] for Matlab [33].

For the NMR contrast analysis, we used the averaged maximum peak of the burst and the signals between bursts as baselines. Calculations were performed using the routine by Gomes et al. [34] which was implemented in Matlab [33]. Preprocessing included the following; Rescaling, which was applied to all data sets before any analysis using the MR vendor’s instructions. Visual inspection of average time series in search for irregularities which were manually removed from the analysis leaving the rest of the time series unaltered. Manual segmentation was used to create a mask to remove cerebral spinal fluid (CSF) contributions. The first 100 of 1000 scans were removed to avoid signal saturation effects. The manual segmentation of the masks was eroded to avoid partial volume effects at the edges.

For the analysis of sleeping pattern, we used a Recurrence Quantification Analysis and a Multifractal Detrended Fluctuation Analysis (for detailed description see Lopez-Perez et al. [35]).

All data graphics were created with Mathematica [36]. Data and source code for analysis are available at www.github.com/Mirandeitor/Entanglement-witnessed-in-the-human-brain.

## IV. DISCUSSION

The aim of this study was to find evidence that brain functions can create entanglement in auxiliary quantum systems.

Thereby, we employed a hybrid MRI sequence which could contain SQC and ZQC, simultaneously. We found that the heart pulsation evoked NMR signal burst with every heartbeat. We were able to show in section II A that the signal contrast originated from spin-spin interactions. Therefore, it is possible that we witnessed quantum entanglement.

However, NMR signals can be altered by many physiological changes. Ultimately, we had to prove that the signal bursts was not a “classical” ZQC.

As mentioned above, classical ZQC have corresponding contrasts in SQC, namely T2* relaxation and diffusion. Both contrasts alter during the heart cycle.

However, T2* changes have shown a different temporal (shifted by more than half of the cycle time in respect to the ZQC signal) and spatial response (higher signal at blood vessel) [37]. The tissue response at around 2% is much lower than during functional activation. In contrast, functional activations showed no significant changes in the ZQC burst signal and only minimal signal increases at the baseline [38]. Therefore, we can conclude that local field-inhomogeneity changes are ineligible as a signal source.

Besides local field-inhomogeneity, ZQC depends on order [39] and rotational symmetries [16, 17] which can be probed with diffusion MRI. The order may correlate with the apparent diffusion coefficient (ADC), while the fractional anisotropy (FA) indicates the rotational symmetry breaking. In praxis, MQC signals are higher at decreased ADC and increased FA.

Nakamura et al. [40] have shown that the temporal changes of the ADC-values are in phase with the intracranial volume change while FA-values show a shift by 180°. Our ZQC signals coincided with the transition phase from the highest to the lowest ADC (and vice versa for the FA). From those results, we can conclude that the theoretical optimum (ADC minimal, FA maximal) for a classical ZQC is outside the time window of the ZQC bursts.

We conclude, that our observation has no corresponding SQC contrast.

Further, the signals surpassed the classical bound by far. For “classical fluids”, the S/N of ZQC compared to the conventional MRI signal (SQC) only reaches up to 0.05 at 4 Tesla, experimentally [41, 42]. Our sequence was suboptimal because we replaced a 90° by a 45° RF-pulse (reduction by factor 2), used a 3 Tesla field, and the evolution time was shorter. Therefore, we can conclude that in combination with the EPI readout, that classical ZQC signals weren’t detectable with our sequence. Even more, in the above argument, we discussed baseline signals. Our observations showed fluctuations which, if translated to a classical ZQC, would then be serval magnitudes higher than the actual baseline ZQC signal.

Although, we found that the evoked bursts disappeared at the magic angle which means they have no SQC component, cardiac pulsation can cause flow and motion effects which we further investigated.

We varied slice thickness and TR as possible sequence parameters, which are sensitive to time-of-flight effects. For the slice thickness, the relative signal did not vary significantly (Fig. 2E), for the repetition time, we found the free induction decay dominating the decline (Fig. 2F). Furthermore, when we varied the blood flow with the help of a CO_2_-challenge (Fig. 4), we found no significant response of the burst signal amplitude.

Further, the fact the signal bursts have no significant SQC component (Fig 2A at the magic angle), we can in principle exclude all SQC contrast mechanism including changes in T_1_ and T_2_ relaxation, line narrowing, or magnetic field shifts.

Above, we have established that conventional MR sequences, be it SQC or MQC, are unable to detect the observed signal bursts. Further, we found that the signal amplitude is above the bound which could classically be reached.

By now, it is clear that the evoked signals can only be observed if the necessary condition, that the magnetisation is highly saturated, is met.

We also considered what we called the sufficient condition above. We found that the timing of the signal bursts coincided the first cluster of the HEP [43]. Like the timing, the signal intensity also showed a similar dependence to conscious awareness in this time window [25, 44].

In another study, López Pérez et al. [35] have shown that the complexity of burst signals correlate with psychological test results in short-term memory. This relation is also known in HEPs.

To our knowledge, both, the direct correlation to conscious awareness and short-term memory, are unreported in classical MRI experiments. It underpins that our findings are from the same origin as HEPs and that there is no classical correlate in MRI.

## V. CONCLUSION

The aim of this study was to show that the brain is non-classical. We assumed that unknown brain functions exist which can mediate entanglement between auxiliary quantum systems. The experimental detection of such an entanglement created by the brain would then be sufficient to prove cerebral non-classicality.

We found experimental evidence that such entanglement creation occurs as part of physiological and cognitive processes.

We argued that the ZQC signals are non-local because (a) ZQC signals are above the classical bound, and (b) the signals have no SQC and MQC[45] correlates. Further, we could confirm that the signals are only detectable in combination with reduced classical signals (necessary condition), and that they resemble HEPs which are below verifiability in conventional MRI (sufficient condition). Our findings disapprove the unproven statement that quantum entanglement or coherence can’t survive in the hot and wet environment of the brain.

Beyond the fundamental question we tried to answer here, we found an undiscovered NMR contrast, which can detect brain activity beyond conventional functional MRI. It may have interesting applications in psychology and medicine.

## DECLARATIONS

### Funding

This research project was funded by Science Foundation Ireland from 2011-15 (SFI-11/RFP.1/NES/3051) and supported by Trinity College Institute of Neuroscience.

### Authors’ contributions

Christian Matthias Kerskens: conceptualisation, methodology (physics), writing (original draft), supervision, funding acquisition.

David López Pérez: methodology (analysis), software, acquisition of data, data curation.

## VI. EXTENDED DATA

**Figure 6.**
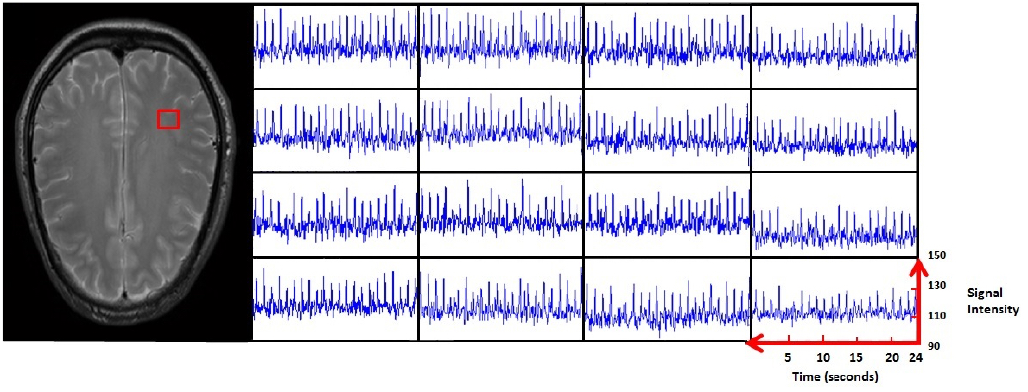
4x4 voxel matrix randomly picked. On the left, the red square shows location in the brain slice. On the right, 16 corresponding signal time courses displaying the local tissue responses over a time period of 24s.

**Figure 7.**
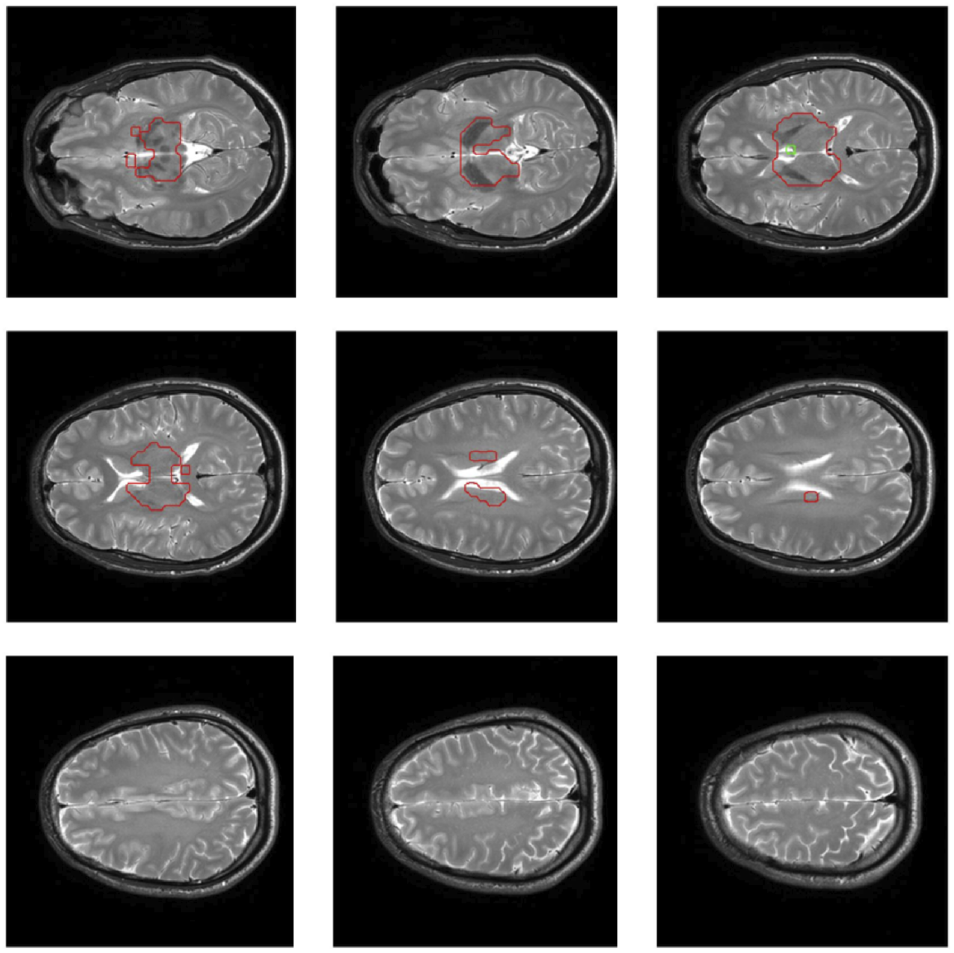
9 Anatomical slices which correspond to the positioning of the EPI time series. Tissue surrounded by red drawing showed no ZQC bursts.

**Figure 8.**
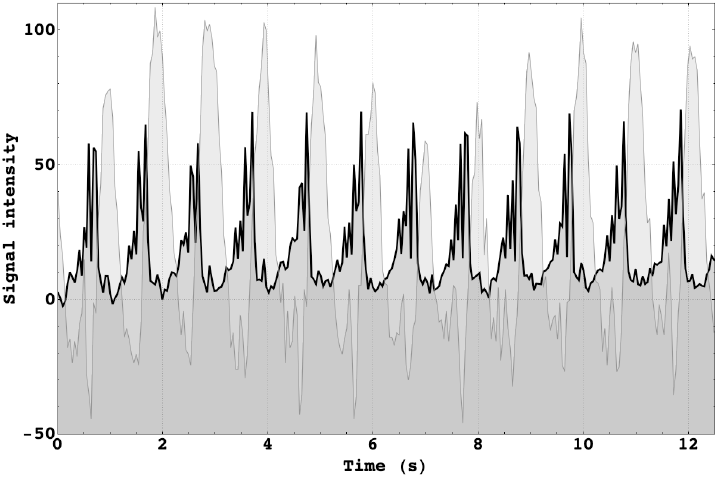
Whole-slice averaged signal time course (black line) which was selected by a mask over 12 heart cycles. Signal of the Superior sagittal sinus (grey line) as reference time frame demonstrates the instant breakdown of quantum coherence with the beginning outflow.

**Figure 9.**
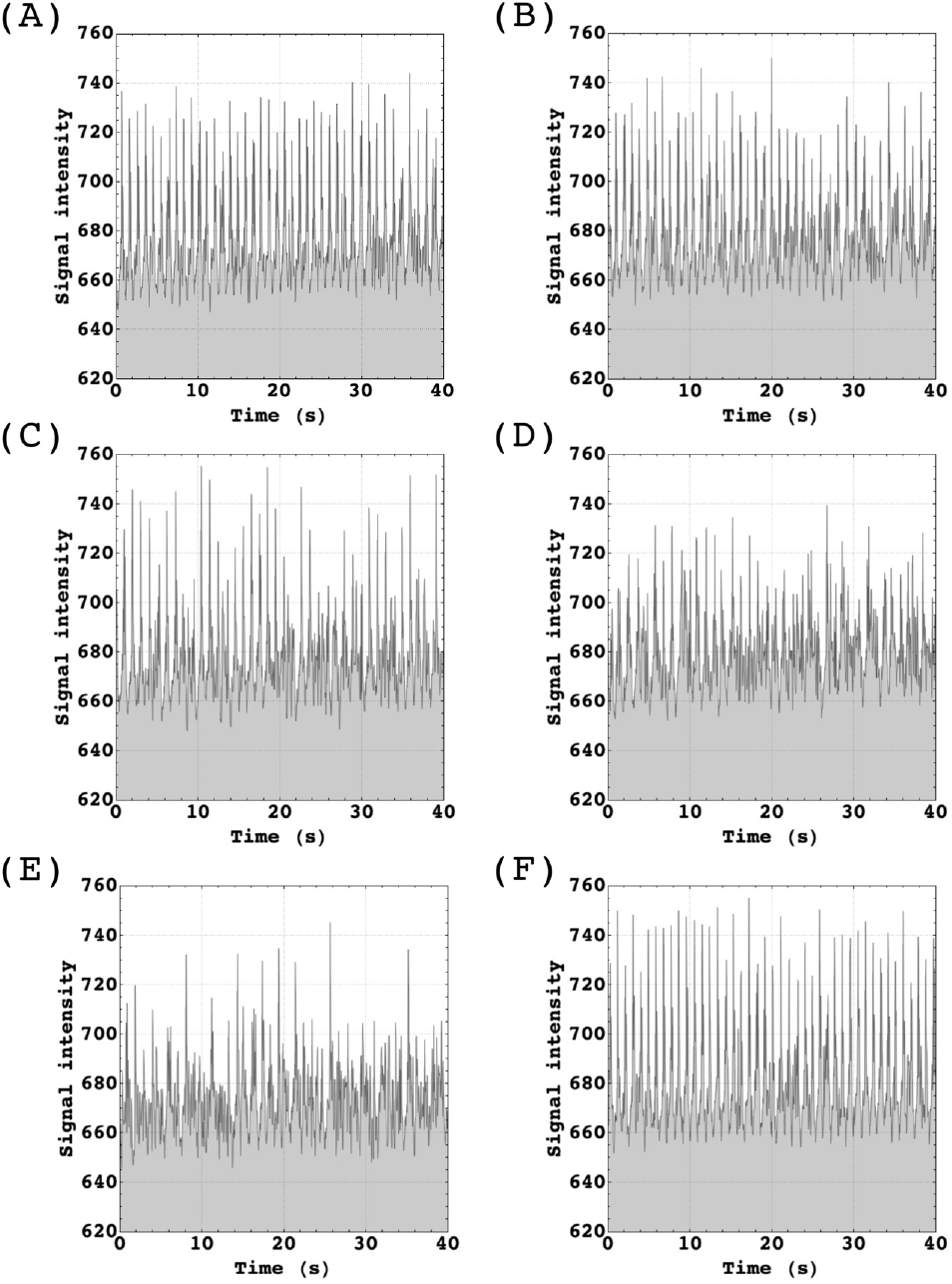
Case study: ZQC burst pattern observed in participant who had reported falling asleep. Starting point of time series at **(A)** 16:26:29 **(B)** 16:29:47 **(C)** 16:30:54 **(D)** 16:34:13 **(E)** 16:37:32 **(F)** 16:40:49 (awake, subject communicated with radiographer before scan).

**Figure 10.**
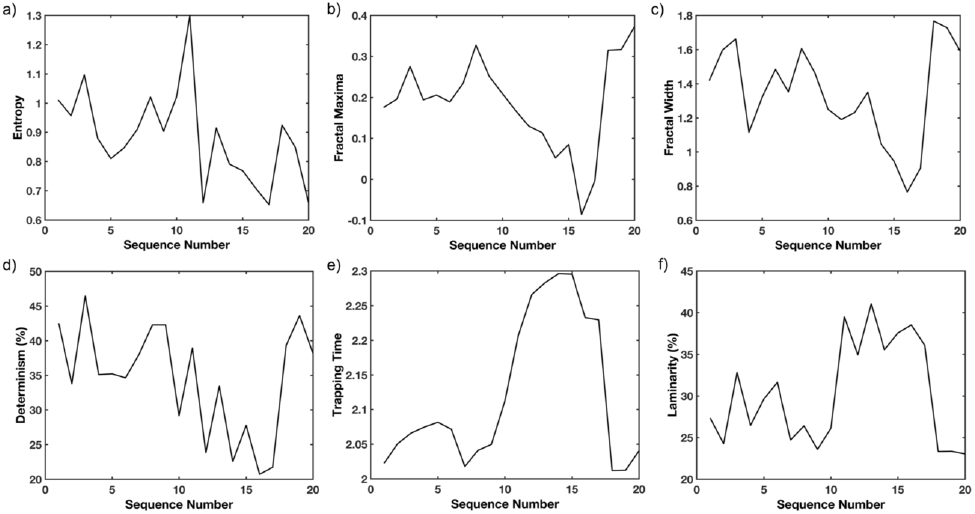
Case study: Results of a Recurrence Quantication Analysis and a Multifractal Detrended Fluctuation Analysis using 20 time periods a 45s over a total time period of 21 minutes. **(a)** Entropy (Ent) is computed as the Shannon entropy of the distribution of the repeating pattern of the system. If a signal has high entropy it exhibits diversity in short and long duration periodicities. **(b-c)** The multifractal spectrum identifies the deviations in fractal structure within time periods with large and small fluctuations. **(d)** Determinism (DET) represents a measure that quantifies repeating patterns in a system and it is a measure of its predictability. Regular, periodic signals, such as sine waves, will have higher DET values, while uncorrelated time series will cause low DET. **(e)** Trapping Time (TT) represents the average time the system remains in a given state and it is a measure of the stability of the system. **(f)** Laminarity (Lam) determines the frequency of transitions from one state to another, without describing the length of these transition phases. It indexes the general level of persistence in some particular state of one of the time-series.

## Notes

https://www.authorea.com/users/270054/articles/393107-cardiac-evoked-long-range-quantum-entanglement-in-the-conscious-brain-supplementary-material

### Competing Interest Statement

The authors have declared no competing interest.

### Summary of Updates

We extended the introduction and discussion substantially.

